# Pulmonary granuloma formation during latent *Cryptococcus neoformans* infection in C3HeB/FeJ mice involves progression through three immunological phases

**DOI:** 10.1101/2024.12.03.626680

**Authors:** Jovany J. Betancourt, Minna Ding, J. Marina Yoder, Issa Mutyaba, Hannah M. Atkins, Gabriella de la Cruz, David B. Meya, Kirsten Nielsen

**Affiliations:** Department of Microbiology and Immunology, University of Minnesota, Minneapolis, MN 55455; Department of Biomedical Sciences and Pathobiology, Virginia Tech University, Blacksburg, VA 24060; School of Medicine, University of North Carolina Chapel Hill, Chapel Hill, NC 27599; Pathology Services Core, University of North Carolina Chapel Hill, Chapel Hill, NC 27599; College of Health Sciences, Makerere University, Kampala, Uganda

## Abstract

*Cryptococcus neoformans* is a fungal pathogen that can cause lethal disease in immunocompromised patients. Immunocompetent host immune responses, such as formation of pulmonary granulomas, control the infection and prevent disseminated disease. Little is known about the immunological conditions establishing the latent infection granuloma in the lungs. To investigate this, we performed an analysis of pulmonary immune cell populations, cytokine changes, and granuloma formation during infection with a latent disease-causing clinical isolate in C3HeB/FeJ mice over 360 days. We found that latently infected mice progress through three phases of granuloma formation where different immune profiles dominate: an early phase characterized by eosinophilia, high IL-4/IL-13, and *C. neoformans* proliferation in the lungs; an intermediate phase characterized by multinucleated giant cell formation, high IL-1α/IFNγ, granuloma expansion, and increased blood antigen levels; and a late phase characterized by a significant expansion of T cells, granuloma condensation, and decreases in lung fungal burden and blood antigen levels. These findings highlight a complex series of immune changes that occur during the establishment of granulomas that control *C. neoformans* in the lungs and lay the foundation for studies to identify critical beneficial immune responses to Cryptococcus infections.

**IMPORTANCE:** *Cryptococcus neoformans* is a fungal pathogen that disseminates from the lungs to the brain to cause fatal disease. Latent C*. neoformans* infection in the lungs is controlled by organized collections of immune cells called granulomas. The formation and structure of Cryptococcus granulomas are poorly understood due to inconsistent human pathology results and disagreement between necrotic granuloma-forming rat models and non-necrotic granuloma-forming mouse models. To overcome this, we investigated granuloma formation during latent *C. neoformans* infection in the C3HeB/FeJ mouse strain which forms necrotic lung granulomas in response to other pathogens. We found that latent *C. neoformans* granuloma formation progresses through phases that we described as early, intermediate, and late with different immune response profiles and granulomatous characteristics. Ultimately, we show that C3HeB/FeJ mice latently infected with *C. neoformans* form non-necrotic granulomas and could provide a novel mouse model to investigate host immune response profiles.

## INTRODUCTION

Pulmonary granulomas are highly variable structures formed by innate and adaptive immune cells surrounding a persistent or difficult to clear agent ^1^. The ability to produce effective pulmonary granulomas is a critical defense mechanism against the systemic dissemination of respiratory pathogens. Cryptococcal meningitis is a fatal disease that accounts for 20% of global HIV-related mortality and is caused by dissemination of the fungal pathogen *Cryptococcus neoformans* from the lungs ^2^. In immunocompetent human *C. neoformans* infections, granuloma formation contains the infection to the lungs and prevents dissemination to the brain ^1,4^. Immunosuppression associated with HIV, cancer, organ transplant therapies, or genetic abnormalities is thought to disrupt the normal granuloma structure and allow for fungal dissemination ^1,5,6^. Indeed, histological analysis of the lungs of immunocompetent and immunosuppressed patients with pulmonary cryptococcal infections showed substantial differences in granuloma structure ^6,7^. However, little is known about how granulomas form during latent infection prior to immunosuppression due to a lack of concise human data.

Analysis of human cryptococcal pulmonary granulomas found necrotic, non-necrotic, and “gelatinous” forms of granulomatous lesions with various inflammatory severities ^1,6,8,9^. Interpreting these findings to predict optimal granuloma structure to control cryptococcal infection is difficult given patients’ complex clinical histories as well as historical issues in identifying the causative Cryptococcus species. Alternatively, *in vivo* models can be used to fill the gap on how granulomas form in immunocompetent hosts during latent infection ^5,10,11^. To accomplish this, it is important to use a model where 1) the *C. neoformans* strain produces a clinically relevant latent infection and 2) the host can produce a human-like granuloma.

Inducing latent cryptococcal infection can be done using patient-derived clinical isolates ^5,11^. While infections using the laboratory reference strain KN99α are known to cause diffuse lung disease that rapidly disseminates and causes fatal cryptococcal meningitis ^12^, we have shown that using the clinical isolate UgCl223, derived from a patient that survived their meningitis, produces a latent infection in immunocompetent mice and a lethal infection in immunosuppressed mice ^5,11^. However, our previous study with UgCl223 was performed in C57Bl/6J mice which are known to be unable to produce necrotic granulomas ^5,13^, limiting the host’s ability to capture the range of human-like granulomatous phenotypes. Therefore, we sought a mouse background capable of also forming necrotic granulomas during pulmonary infection.

C3HeB/FeJ mice form large, necrotic pulmonary granulomas in response to lung infections and are used extensively to study Mycobacterium disease ^13–15^. C3HeB/FeJ mice express the ‘susceptible’ allele of the super susceptibility to tuberculosis 1 (sst1S) locus which influences several immunoregulatory genes ^13,15^. Specifically, sst1S contains deficiencies in the speckled-protein SP140 which negatively regulates type 1 interferon (IFN1) expression and has been found to contribute to the C3HeB/FeJ necrotic granuloma phenotype ^16,17^. It is unknown what granulomatous phenotypes C3HeB/FeJ mice display during pulmonary fungal infections.

In this study, we infected C3HeB/FeJ mice with the latent disease-causing clinical isolate UgCl223 ^5^ to model the immune response involved in forming pulmonary granulomas. Using this model, we identified three phases of granuloma formation comprising different cellular, cytokine, and histological profiles. We observed the formation of non-necrotic, mature granulomas composed of phagocytes, neutrophils, and CD4+ T cells. Further, we found that the cryptococcal C3HeB/FeJ granulomatous immune response was not due to SP140 deficiency, unlike in *M. tuberculosis*. These data show that latent *C. neoformans* infection with a wild-type clinical isolate induces non-necrotic pulmonary granuloma formation in mice and that waves of different immune response profiles precede the formation of mature granulomas.

## MATERIALS AND METHODS

### Ethics Statement

Animal experiments were done in accordance with the Animal Welfare Act, United States federal law, and National Institutes of Health guidelines. Mice were handled in accordance with guidelines defined by the University of Minnesota Animal Care and Use Committee (IACUC) under protocols 1908A37344, 2207A40205, and 2104A39016.

### Mouse Infections

C3HeB/FeJ (Strain #000658, Jackson Laboratories, Bar Harbor, ME), C57Bl/6J (Strain #000664, Jackson Laboratories, Bar Harbor, ME), or SP140^-/-^ C57Bl/6J (SP140^-/-^ B6) ^17^ mice were infected intranasally with 10^3^ cells of the *C. neoformans* serotype A clinical isolate UgCI223 ^5,11^ or the *C. neoformans* serotype A laboratory reference strain KN99α ^18^ to generate latent or lethal infections, respectively, and then monitored for up to 360 days post-infection (DPI). Mice used for determining disease outcomes, histology, and cytokines were sacrificed at 0, 10, 20, 30, 60, 90, 120, 180, 240, 300, and 360 DPI using CO_2_ euthanasia. Mice used for flow cytometry were sacrificed at 0, 14, 21, 30, 59, 91, 150, 210, 270, and 360 DPI. For survival curves, 8-10 mice were infected as described above and monitored for *C. neoformans* disease. Mice were sacrificed when they exhibited neurological symptoms and/or lost 20% of initial weight.

### Cryptococcal Antigen Testing

Blood was collected prior to sacrifice in tubes containing 100 mM EDTA, and antigenemia was assessed using cryptococcal antigen (CrAg) lateral flow assay (LFA) dipsticks (IMMY, Norman, OK) following the manufacturer’s instructions. Briefly, blood was diluted 1:5 and doubled until the highest dilution that yielded a positive result was identified or a maximum titer of 1:2560 was reached. Samples positive at ≥1:640 were considered evidence of breakthrough infection and excluded from analysis.

### Tissue Fungal Burden

Lung and brain tissue were excised, submerged in phosphate buffered saline (PBS), and homogenized. Half of the lung homogenate was saved for cytokine analysis. The remaining tissue homogenate was used for fungal burden enumeration via serial dilution plating on yeast peptone dextrose (YPD) plates with colony forming units (CFUs) counted after 48 hours of incubation at 30°C.

### Cytokine Analysis

Lung homogenate was mixed with proteinase inhibitor (cOmplete EDTA-free, Roche, Indianapolis, IN), centrifuged, supernatant frozen in liquid nitrogen, and stored at -80°C until analyzed. ProcartaPlex kits (Thermo Scientific, Waltham, MA) were used to quantify cytokine abundances following the manufacturer’s instructions, using a Luminex Multiplex Immunoassay System (Thermo Scientific, Waltham, MA). Cytokines analyzed were IFNα, IFNγ, IL-2, IL-1β, IL-4, IL-6, IL-10, IL-12p70, IL-13, IL-17, GM-CSF, TNFα, eotaxin, MIP-1α, MIP-1β, IP-10, and RANTES.

### Flow Cytometric Analysis

Lung tissue was collected, digested, and purified for CD45+ cells using the StemCell Easy Eights CD45 Positive Selection kit (StemCell Technologies, Vancouver, Canada), as described previously ^5^. CD45+ cells were stained for surface markers as per manufacturer instructions. Samples were analyzed using a BD LSRFortessa X-20 (Becton Dickinson, Franklin Lakes, NJ). Total cell counts and proportions were analyzed using FlowJo software (Ver 10.10, Becton Dickinson, Franklin Lakes, NJ).

A 14-color bulk CD45+ cell panel was performed to identify B cells, T cells, Ly6C+ monocytes, macrophages (alveolar, interstitial, and exudative), CD11b+ and CD11b-dendritic cells (DCs), natural killer (NK) cells, eosinophils, and neutrophils as previously described (Ding et al., 2022). The bulk panel antibodies used were TCRβ (H57-597, PE, Biolegend), CD11b (M1/70, APC, BD Biosciences), CD11c (HL3, BV786, BD Biosciences), Ly6C (Hk1.4, PerCp-Cy5.5, Invitrogen), MHCII (M5/114.15.2, BV605, Biolegend), CD24 (M1/69, BV711, BD Biosciences), CD64 (X54-5/7.1, BV421, Biolegend), SiglecF (E50-2440, PE-CF594, BD Biosciences), Ly6G (1A8, AF700, BD Biosciences), B220 (RA3-6B2, BV650, Biolegend), NK1.1 (PK136, AF488, Biolegend), CD45 (30-F11, BUV805, Invitrogen), intravenous CD45 to identify circulating immune cells (30-F11, BUV395, BD Biosciences), and Live/Dead (APC-eFluor780, Invitrogen).

A 15-color T cell panel was performed to identify CD4+ and CD8+ naïve, effector, regulatory (Treg), and tissue resident memory (Trm) cells. The T cell panel antibodies used were TCRγ/δ (GL3, PE, Biolegend), PD-1 (J43, APC, Invitrogen), NK1.1 (PK136, BV786, BD Biosciences), CD69 (H1.2F3, PerCp-Cy5.5, Invitrogen), CXCR3 (CXCR3-173, BV605, Biolegend), CD8 (53-6.7, BV711, Invitrogen), CD62L (MEL-14, BV421, Biolegend), CD44 (IM7, PE-CF594, BD Biosciences), TCRβ (H57-597, AF700, Biolegend), CD25 (PC61, BV650, Biolegend), CD103 (RMST2-2, AF488, Invitrogen), CD4 (GK1.5, BUV805, BD Biosciences), B220 (RA3-6B2, APC-eFluor780, Invitrogen), CD11c (N418, APC-eFluor780, Invitrogen), F4/80 (BM8, APC-eFluor780, Invitrogen), intravenous CD45 to identify circulating immune cells (30-F11, BUV395, BD Biosciences), and Live/Dead (APC-eFluor780, Invitrogen).

### Histology

Whole lungs and brain were excised, fixed in 10% neutral-buffered formalin for 72-120 hours, and transferred into 70% ethanol for storage. Fixed organs were paraffin-embedded, sectioned, mounted on slides, and stained at the Pathology Services Core at the University of North Carolina-Chapel Hill (PSC). Slide-mounted sections were stained with hematoxylin and eosin (H&E) for visualization or with one of two fluorescent-linked antibody panels for localization. The antibodies used for immunofluorescent panel 1 were F4/80 (Cy5, 1:2000), CD4 (Cy3, 1:150), and CD68 (AF488, 1:200) and panel 2 were EPX (Cy5, 1:700), Ly6G (Cy3, 1:1000), and CD68 (AF488, 1:200). Slides were digitalized using the Aperio ScanScope FL (Aperio Technologies Inc). The digital images were captured in each channel by 20x objective (0.468 μm/pixel resolution) using line-scan camera technology (U.S. Patent 6,711,283). The adjacent 1 mm stripes captured across the entire slide were aligned into a contiguous digital image by an image composer. Images were archived in PSC’s eSlide Manager database (Leica Biosystems). Digital image visualization was performed using Aperio ImageScope (Ver. 12.4.6, Leica, Wetzlar, Germany).

### Statistical Analysis

Survival curves were analyzed for significance using the Kaplan-Meier estimate. CFUs and LFA titers were statistically evaluated using one-way ANOVA with Tukey’s post-hoc correction for multiple comparisons. Cytokines and immune cell count and proportions were statistically evaluated using one-way ANOVA and compared to the uninfected control. All statistical analyses were performed using GraphPad Prism software (Ver. 10.2.0, Dotmatics, Boston, Massachusetts).

## RESULTS

### C3HeB/FeJ Mice with Latent *Cryptococcus neoformans* Infections Exhibit Long-Term Survival

A model of cryptococcal granuloma formation is needed to understand effective host immune responses that control latent *C. neoformans* infection. We previously developed a latent model of *C. neoformans* using C57Bl/6J mice and the clinical isolate UgCl223 ^5^. While granulomas were observed, they had disorganized structures and >20% exhibited fatal reactivation/breakthrough, making it challenging to study the progression of the granulomatous immune response. In contrast, C3HeB/FeJ mice are known to form organized, necrotic pulmonary granulomas when infected with intracellular pathogens, such as *M. tuberculosis* and *M. bovine,* that better recapitulate human granulomas ^13,15,17^. Like these pathogens, *C. neoformans* is known to form long-term, stable pulmonary granulomas in humans ^1,10^. To determine whether the C3HeB/FeJ granulomatous response would better mimic human latent *C. neoformans* infection, C3HeB/FeJ mice were infected with 1x10^3^ cells of the latent UgCl223 clinical isolate or the lethal laboratory reference strain KN99α and sacrificed at designated timepoints over 360 days. C3HeB/FeJ mice infected with lethal KN99α had a median survival of 20 days while mice infected with latent UgCl223 survived indefinitely (Fig. 1A). As in the C57Bl/6J latent infection model, breakthrough lethal infections occurred in the C3HeB/FeJ mice between 60-90 DPI, but the proportion of breakthrough infections was lower with only 12.5% breakthrough observed in the C3HeB/FeJ mice compared to >20% in the C57Bl/6J mouse background. In C3HeB/FeJ mice, lung fungal burden peaked at 20 DPI (Fig. 1B), and no significant increase in brain fungal burden was observed (Fig. 1C).

**Figure 1:**
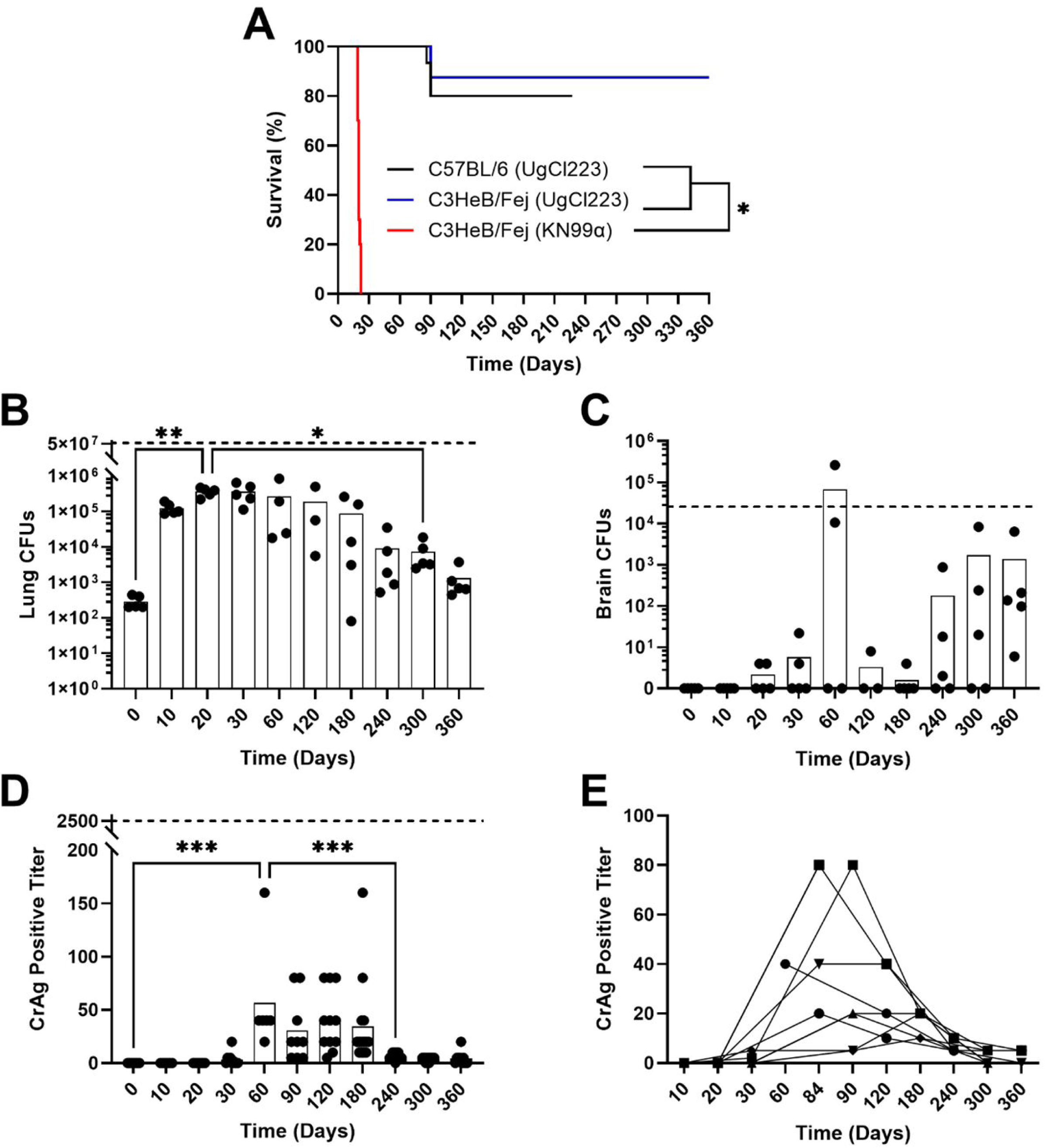
Disease outcomes of latent cryptococcal infection in C3HeB/FeJ mice. A) Kaplan-Meier survival curves for C3HeB/FeJ and WT-B6 mice infected with latent UgCl223 or KN99α. Tissue fungal burdens for B) lungs and C) brain. Dashed-line indicates the mean CFUs for KN99α infection at 20 DPI. D) CrAg LFA maximum positive titers for sacrificed and survival cohort mice. Dashed-line indicates the mean LFA titer for KN99α infection at 20 DPI. E) LFA titers for survival cohort mice over time with lines connecting titers collected from the same mouse. B, C, and D were analyzed via one-way ANOVA with Tukey’s post-hoc.* = p-value <0.05, *** = p-value <0.001.

Cryptococcal antigen lateral flow assays (CrAg LFAs) are used clinically to diagnose disseminated cryptococcosis (i.e., antigenemia) and identify patients at-risk for fatal cryptococcal meningitis ^19^. To determine the presence and severity of dissemination in the mouse model, CrAg LFAs were used to determine blood antigen levels. The highest antigenemia levels in the latent UgCl223 infections were observed during the 60-90 DPI range, concurrently with disease breakthrough, before dropping to near negative titers (Fig. 1D-E). Importantly, antigen levels in the latent UgCl223 infections were substantially lower than those observed in the lethal C3HeB/FeJ KN99α infections at 14 DPI, where blood titers were positive at the 1:2560 dilution (lethal KN99α levels are indicated as dashed lines in Fig. 1B-D).

Taken together, these data show that C3HeB/FeJ mice infected with UgCl223 establish latent infections, survive indefinitely, have lower levels of lethal breakthrough infections than C57Bl/6J mice and have minimal evidence of extrapulmonary dissemination.

### Cryptococcus Pulmonary Granuloma Structure Changes Over Time

We next sought to determine whether the distinctive C3HeB/FeJ mouse immune response to latent cryptococcal infection affected granuloma formation. We investigated the histopathological progression of latent Cryptococcus disease over 360 days in C3HeB/FeJ mice infected with UgCl223 (Fig. 2). Characteristics such as the presence of *C. neoformans* cells, immune cell populations, and changes in lung architecture were used to determine the severity of infection, visualize immune responses, and follow granuloma formation. At 10 DPI lung structure was largely normal with no signs of an active immune response despite the observation of several *C. neoformans* cells in the alveolar spaces. From 20-30 DPI, we observed phagocytes actively phagocytosing *C. neoformans* and the recruitment of polymorphonuclear cells (PMNs) such as eosinophils and neutrophils. The pulmonary lesions were localized, with normal alveolar structure in areas without inflammation. At 60 DPI large immature granulomas were observed containing lymphocytes and multinucleated giant cells (MNGCs) with phagocytosed *C. neoformans*. Importantly, these granulomas did not have necrotic cores, with the tissue outside of the granulomas exhibiting near-normal histology (some compression of alveoli is seen around large granulomas). Between 90-120 DPI, the severity of pulmonary infection lessened, with fewer *C. neoformans* cells observed within the granulomas. Lymphocytes and MNGCs remained the most prominent immune cell types. From 180-360 DPI, the lymphocyte density increased considerably. Mature granulomas containing lymphocytes surrounding a core of phagocytes engulfing *C. neoformans* were observed starting at 180 DPI (Fig. 2 180 DPI). The surrounding pulmonary tissue was appreciably unaffected with normal alveolar septa thickness, appropriately sized small airways, and no appreciable scarring.

**Figure 2:**
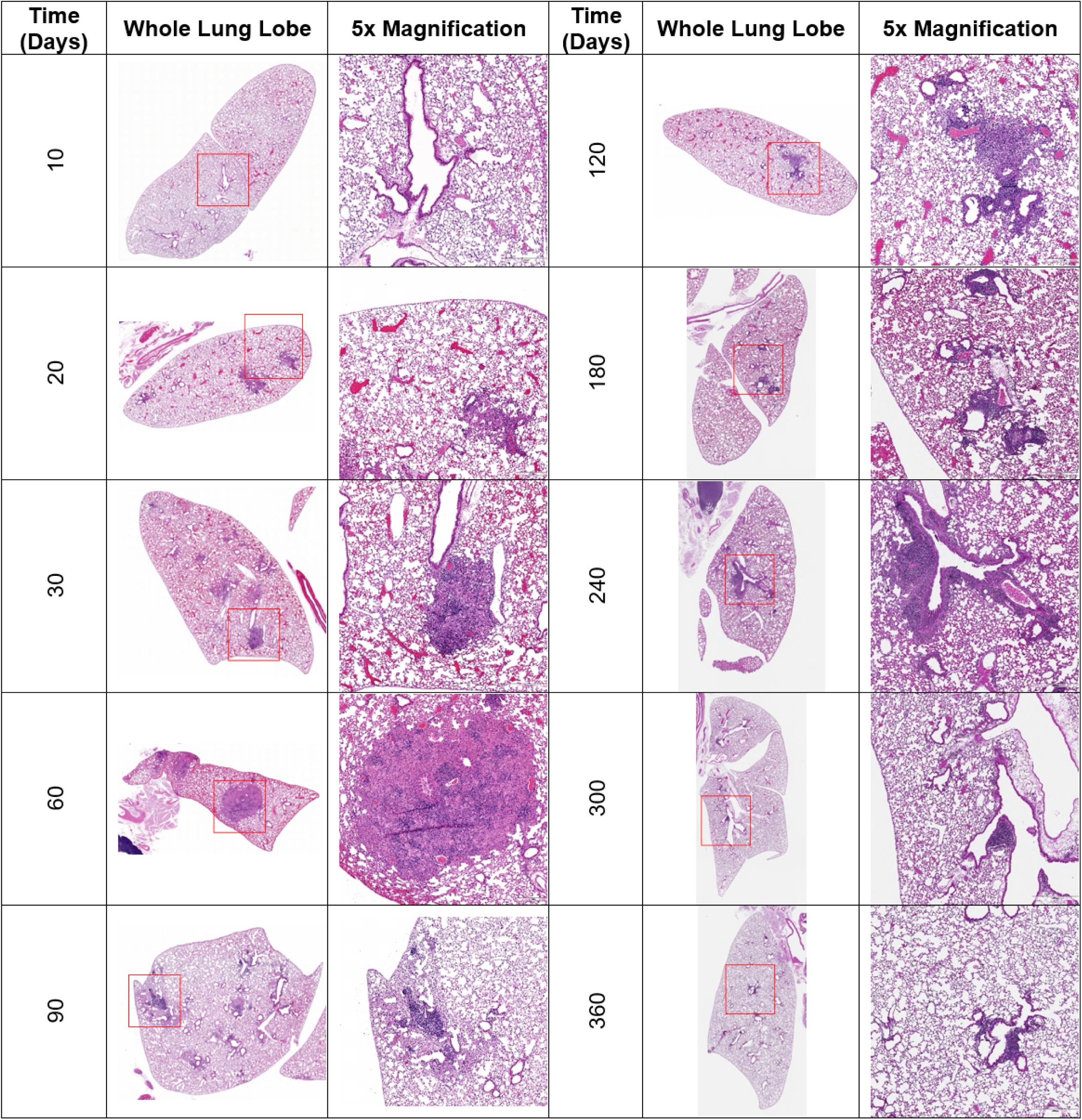
C3HeB/FeJ mice form non-necrotic pulmonary granulomas composed primarily of CD68+ macrophages, neutrophils, and CD4+ T cells. Lungs were collected from infected C3HeB/FeJ mice at predetermined timepoints over 360 days and stained with hematoxylin and eosin. Number of mice lungs imaged per timepoint (n) = 5. Representative images of whole lung and a 5x magnified region for each timepoint are shown.

These histopathological findings show that even in a mouse background known to produce necrotic pulmonary granulomas, latent Cryptococcus infection produced mature, non-necrotic granulomas, with the earliest mature granuloma found at 180 DPI. Furthermore, these findings show that latent infection induces a complex series of histopathological changes that predate the establishment of the mature granuloma.

### C3HeB/FeJ Mice Produce a Different Immune Response to Latent Cryptococcus Infection Compared to C57Bl/6J Mice

Differences in pulmonary cell populations could impact the formation and structure of granulomas ^1^. To determine whether C3HeB/FeJ and C57Bl/6J mice exhibited differences in pulmonary cellular profiles during latent *C. neoformans* infection, we analyzed lung homogenates using flow cytometry to quantify eleven immune cell types. We measured the abundance of T cells, B cells, eosinophils, neutrophils, natural killer (NK) cells, CD11b+ dendritic cells (DCs), CD11b-DCs, Ly6C+ monocytes, alveolar macrophages (AMacs), interstitial macrophages (IMacs), and exudative macrophages (ExMacs) (Fig. 3). Our results show that C3HeB/FeJ mouse lungs contain significantly less total CD45+ immune cells compared to C57Bl/6J mice at 30 days (Fig 3 CD45+ Cell Counts). Analysis of cell proportions across the mouse backgrounds showed the C3HeB/FeJ mice had a lower proportion of Ly6C+ monocytes but significantly higher proportions of eosinophils, CD11b+ DCs, and neutrophils than observed in the C57Bl/6J mice at 30 DPI (Fig 3).

**Figure 3:**
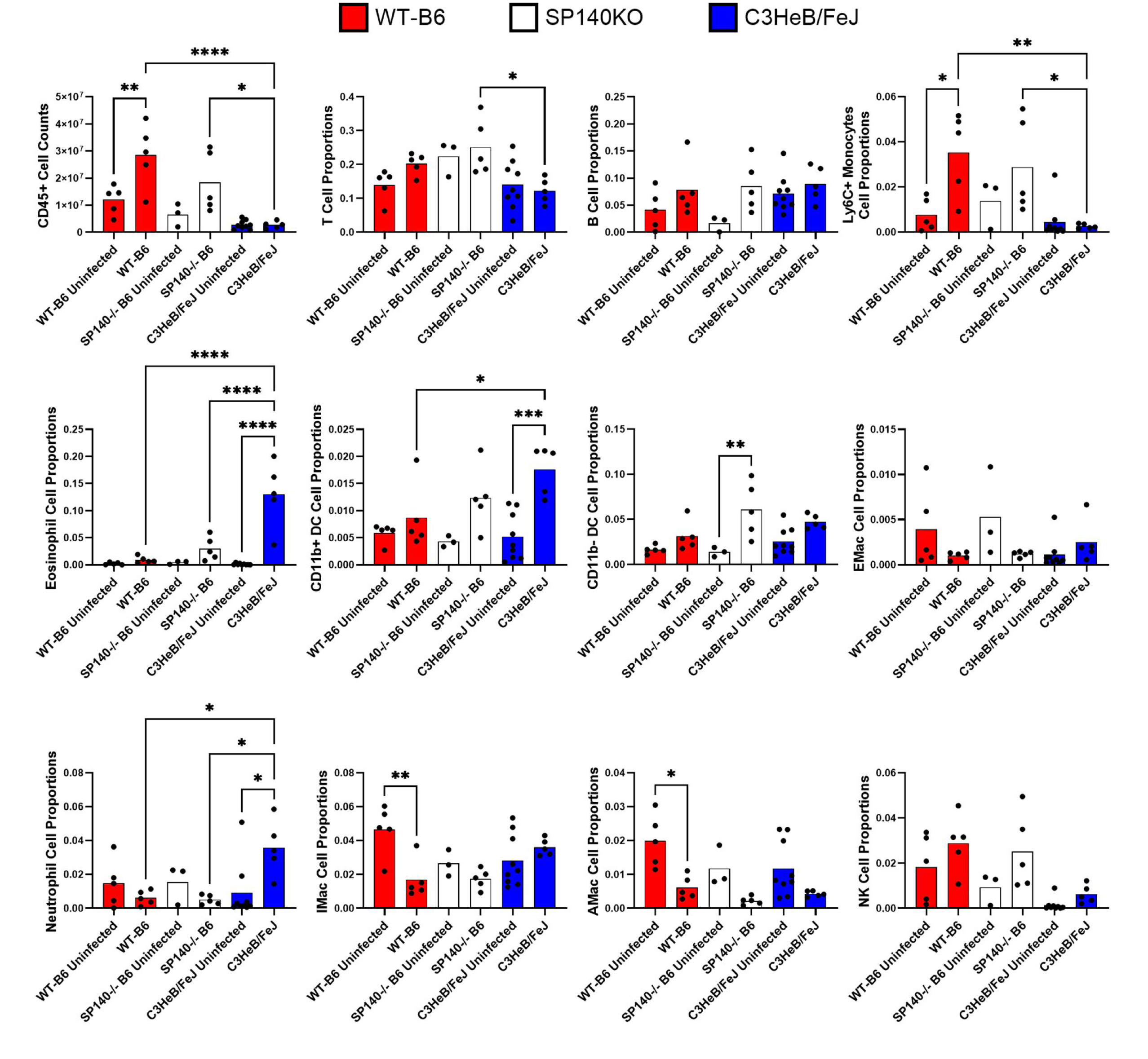
SP140KO mice do not replicate the immune response seen in C3HeB/FeJ mice during cryptococcal infection. Lungs of UgCl223-infected C3HeB/FeJ, wild-type C57Bl/6 (WT-B6), and SP140^-/-^ C57Bl/6J (SP140-/-B6) were collected at 30 DPI and analyzed by flow cytometry to enumerate eleven different CD45+ cell types. The top-left graph depicts total CD45+ cell counts while the rest of the graphs depict that cell type’s proportion of all CD45+ cells. Groups were analyzed via one-way ANOVA with Tukey’s post-hoc. Only biologically relevant pairwise comparisons are shown (i.e., comparisons within the same mouse background or comparisons of different mouse backgrounds of same infection status). * = p-value <0.05, ** = p-value <0.01, **** = p-value <0.0001.

These data show that C3HeB/FeJ mice exhibit a unique response to latent Cryptococcus infection characterized by a distinct population of CD45+ immune cells in the lungs compared to the C57Bl/6J model. C3HeB/FeJ mice recruit less CD45+ immune cells with a different innate immune cell population composition.

### SP140 Deficiency Does Not Replicate the C3HeB/FeJ Immune Response to Latent Cryptococcus Infection

Previous studies with C3HeB/FeJ mice identified the genetic locus sst1S as being the main contributor to the necrotic granuloma structure during *M. tuberculosis* infection ^13,15,17^. Further studies of genes within this locus revealed a deficiency in the speckled-protein family member SP140 allele, which functions as a negative regulator of IFN1. Using SP140^-/-^ C57Bl/6J mice, Ji et al. demonstrated that deficiency in SP140 significantly increases the abundance of IFN1 and is important for producing necrotic pulmonary granulomas during *M. tuberculosis* infection ^17^.

To determine if SP140 deficiency influences the granulomatous response to latent Cryptococcus infection in an IFN1-dependent manner, SP140^-/-^ C57Bl/6J mice were infected with UgCl223 and compared to wild-type C57Bl/6J (WT-B6) and C3HeB/FeJ mice at 30 days post-infection. We found that SP140 deficiency alone in C57Bl/6J mice does not replicate the unique immune response produced by C3HeB/FeJ mice during latent Cryptococcus infection (Fig. 3). Tissue fungal burden was not significantly different across the three mouse backgrounds (Supp. Fig. 1). Interestingly, SP140^-/-^ C57Bl/6J mice did exhibit significant increases in CD11b-DCs during latent Cryptococcus infection when compared to WT-B6 or C3HeB/FeJ mice. There were also several significant differences between the SP140^-/-^ C57Bl/6J and the C3HeB/FeJ mice. *M. tuberculosis* infections of C3HeB/FeJ and SP140^-/-^ C57Bl/6J mice are characterized by robust interferon 1 (IFN1) production ^17^. Surprisingly, while we observed IFN1 production during lethal KN99α infection, we did not observe significant production of IFN1 cytokines during the latent UgCl223 infection (Supp. Fig. 2). These data show that latent Cryptococcus infection in C3HeB/FeJ mice does not induce IFN1 production, that SP140 deficiency does not enhance IFN1 production during latent infection, and that SP140 deficiency in C57Bl/6J mice does not replicate the latent Cryptococcus infection immune response seen in C3HeB/FeJ mice.

### Control of Latent Cryptococcus Infection Involves Progression Through Multiple Phases of Granuloma Formation

Our previous studies using C57Bl/6J mice showed that myeloid and lymphoid immune responses are induced in response to latent infection ^5^. However, the diffuse nature of the C57Bl/6J granuloma made it difficult to track changes in the granulomatous immune response over time. The more focal granulomas observed in C3HeB/FeJ mice allowed us to investigate the granuloma’s cellular and signaling landscape. We analyzed cellular populations and cytokine abundances using flow cytometry and multiplex assays, respectively, to characterize the immune responses generated during C3HeB/FeJ granuloma formation. As described below, these analyses revealed three phases in which different immune cellular and signaling responses dominated. Based on the relative abundance of different cells and cytokines over 360 days, we defined these phases as the early phase (0-30 DPI), intermediate phase (30-180 DPI), and the late phase (150-360 DPI) of granuloma formation.

Cytokines are crucial for cellular signaling during the initiation or termination of immune responses. We analyzed sixteen cytokines known to drive type-1 (IFNγ, IL-2), type-2 (IL-4, IL-13), type-17 (IL-17A), as well as proinflammatory and anti-inflammatory responses (IL-1β, IL-6, IL-10, IL-12p70, IP-10, TNFα, MIP-1α, MIP-1β, eotaxin, RANTES, GM-CSF). These cytokines were clustered based on their relative abundances using Pearson correlation to group cytokines with similar temporal expression patterns (Fig. 4). We observed four major groups of expression patterns: 0-10 DPI contained the non-specific tissue injury cytokines IL-6 and GM-CSF; 20-30 DPI contained the type-2 cytokines IL-4 and IL-13; 30-180 DPI contained a mix of type-1 (IFNγ), type-17 (IL-17A), and proinflammatory cytokines (IL-12p70, MIP-1β, TNFα, IL-2, IL-1β, and MIP-1α); and 240-360 DPI contained a mix of the T cell signaling cytokine RANTES, an anti-inflammatory cytokine IL-10, and the proinflammatory cytokines eotaxin and IP-10. Consistent with these findings, analysis of each cytokine revealed significant changes in abundance over time that coincided with the three phases of the immune response (Supp. Fig. 3).

**Figure 4:**
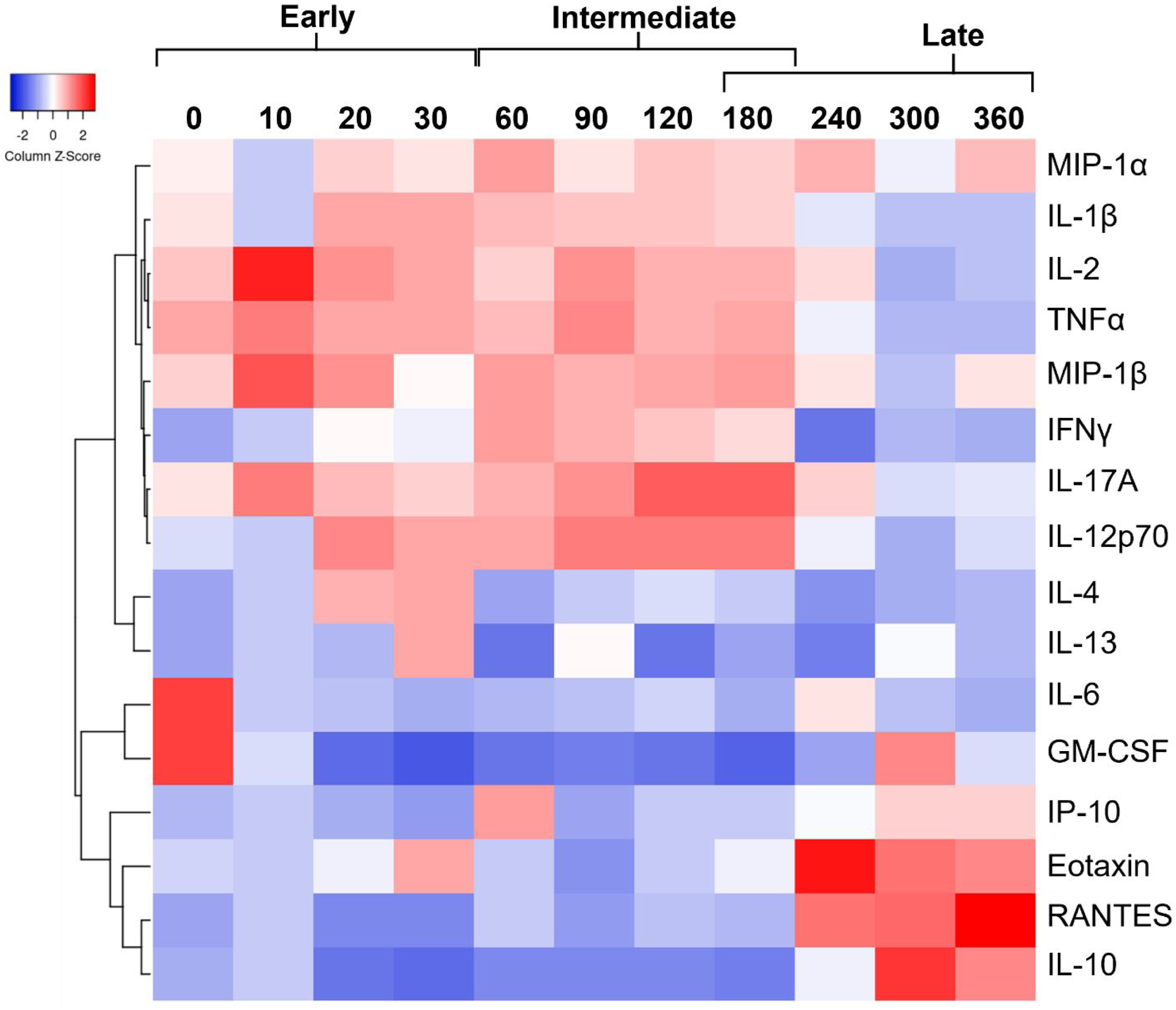
Pulmonary cytokine levels change during latent infection. Pearson-correlation-clustered heatmap of pulmonary cytokine profiles from C3HeB/FeJ mice latently infected with UgCl223 at designated timepoints between 0- and 360-days post-infection. Cytokine levels from lung serum were measured using a ProcartaPlex multiplex immunoassay kit. Early, intermediate, and late indicate phases of granuloma formation. Number of lung serum samples (n) = 3-5 for each column.

Investigating the cellular landscape during pulmonary infection provides insight into the key cell types involved in granuloma formation. We analyzed T cells, B cells, eosinophils, neutrophils, NK cells, CD11b+ DCs, CD11b-DCs, Ly6C+ monocytes, AMacs, IMacs, and ExMacs over the 360-day latent infection. During the UgCl223 infection, there was an initial expansion of myeloid cells in the early phase that retracted during the intermediate and late phases of granuloma formation as lymphocyte populations increased (Fig. 5A). The dominant myeloid cell type transitioned from eosinophils in the early phase to neutrophils and IMacs during the intermediate and late phases. This contrasts with the KN99α infection, where severe myeloid cell expansion occurred instead of the myeloid retraction observed in the latent infection. Immune cell count was consistent with the observed changes in cellular proportions (Fig. 5B, Suppl. Fig. 4). Eosinophils and CD11b+ DCs increased at 21 DPI and then quickly dropped to baseline by 30 DPI. The abundance of CD45+ cells increased by 150 DPI, driven by increases in T cells, B cells, and neutrophils. IMacs and CD11b-DCs reached significant levels by 91 DPI. Interestingly, while the lethal KN99α infection (indicated by the dashed line in the figure) induced a greater total number of immune cells at 20 DPI when compared to the latent UgCl223, we found that mice infected with UgCl223 had more CD11b+ DCs at 20 DPI (Suppl. Fig. 4).

**Figure 5:**
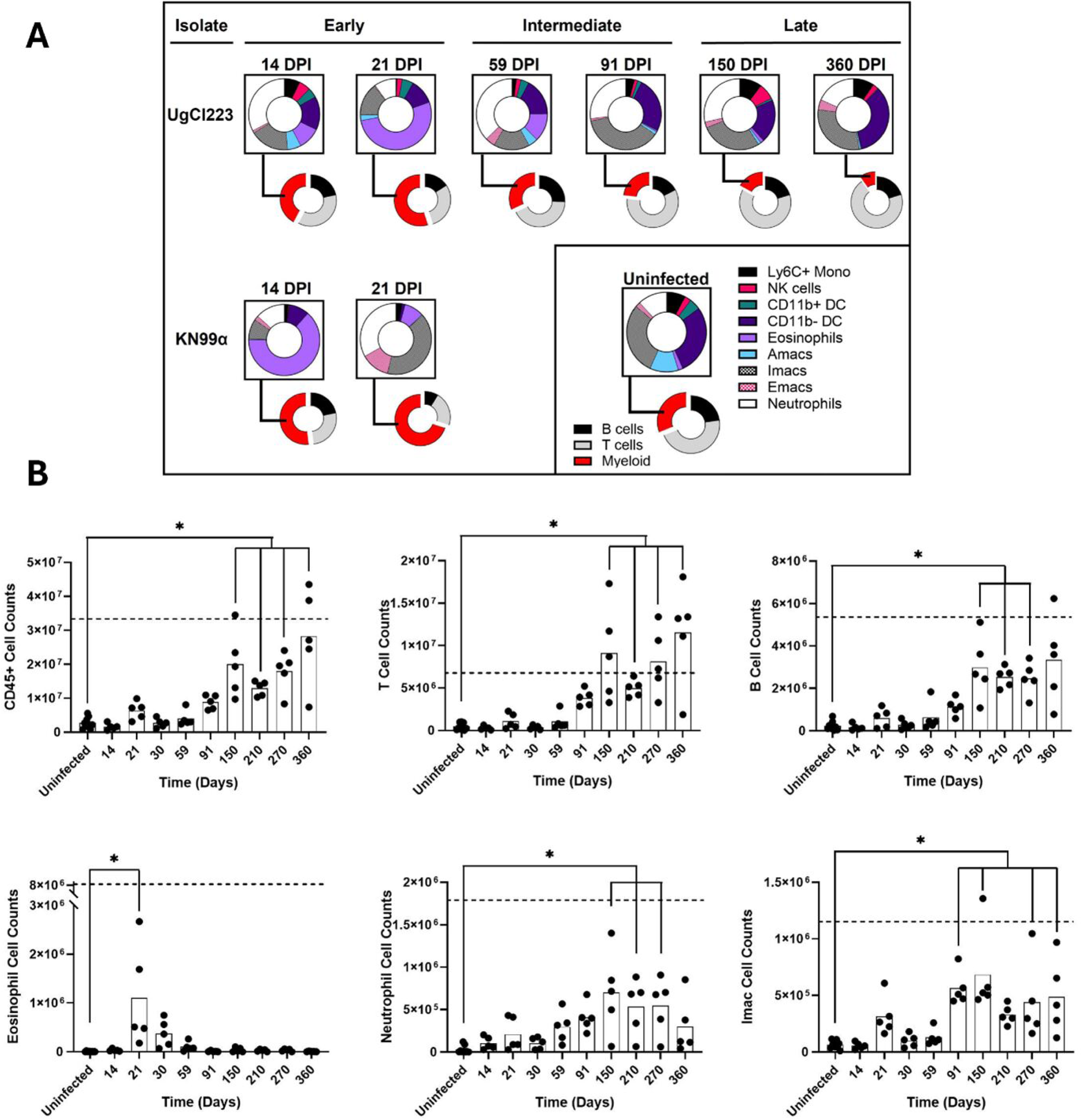
Pulmonary immune cell populations change during latent infection. A) Proportion of pulmonary CD45+ cells in C3HeB/FeJ mice infected with latent UgCl223 or lethal KN99α strains. Representative timepoints from each phase are displayed. B) Total cell counts for CD45+, T cell, B cell, eosinophil, neutrophil, and interstitial macrophages (IMac). Dashed lines represent cell counts for KN99α at 14 DPI. Tissues were collected from UgCl223 infected mice at designated timepoints or KN99α mice at intermediate (14 DPI) or late (21 DPI) time points of the lethal infection. Ordinary one-way ANOVA comparing means to uninfected. * = p-value ≤ 0.05.

T cells consistently provide crucial protection during Cryptococcus infection ^5,20^. To determine changes in T cell subsets during granuloma formation, we analyzed naïve, effector, regulatory, circulating memory, tissue resident memory, and PD1+ exhausted CD4+ and CD8+ T cell subsets. Our results show that both CD4+ and CD8+ T cells experience significant changes in subset populations during latent infection (Fig. 6). Naïve CD4+ T cells increased significantly at 210 and 360 DPI, and naïve CD8+ T cells increased at 210, 270, and 360 DPI. At 21, 150, and 360 DPI there were increases in regulatory and PD1+ CD4+ T cells. CD8+ regulatory T cells increased only at 21 DPI and CD8+ PD1+ T cells increased at 21, 91, 150, and 360 DPI. At 91, 150, and 360 DPI, the effector CD8+ T cells showed increases. Similarly, CD4+ effector T cells increased at 150 and 360 DPI. CD4+ and CD8+ circulating memory and tissue resident memory T cells increased significantly only at 360 DPI. In summation, regulatory T cell populations significantly increased during the early, intermediate, and late phases. Effector T cells increased during the intermediate and late phases. Significant naïve and tissue resident memory T cell expansion only occurred during the late phase.

**Figure 6:**
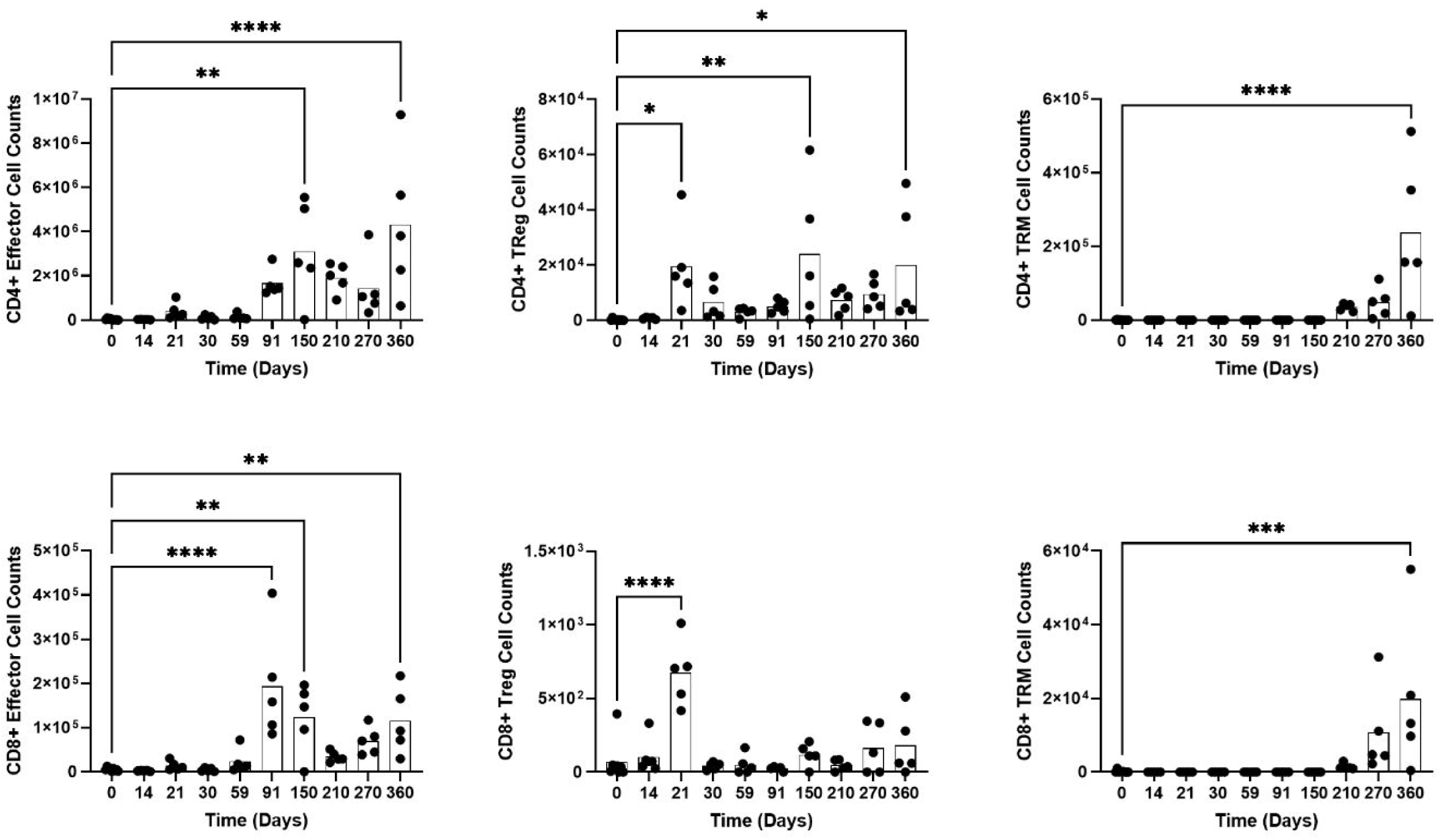
Pulmonary CD4+ and CD8+ T cell populations experience differential expansion. Total cell counts for CD4+ and CD8+ effector, regulatory (Treg), and tissue-resident memory (TRM) cells. Lungs were collected from UgCl223 infected mice at designated timepoints, homogenized, and analyzed using a T cell-specific flow cytometry panel. Ordinary one-way ANOVA comparing means to 0 DPI. * = p-value <0.05, ** = p-value <0.01, *** = p-value < 0.001, **** = p-value <0.0001.

### C3HeB/FeJ Cryptococcus Granulomas Develop a Highly Organized Structure

Our analyses identified eosinophils, neutrophils, CD68+ macrophages, and CD4+ T cell populations dramatically change in abundance across time. We sought to localize these cells during granuloma formation to determine their spatial orientation in the Cryptococcus granuloma. To accomplish this, we utilized immunofluorescence staining (IF) to stain primary markers and localized their presence in pulmonary tissue. We targeted macrophages by F4/80 and CD68, CD4+ T cells by CD4, neutrophils by Ly6G, and eosinophils by EPX. Our data reveal that the presence of these cells within and around the granulomas change over time (Fig. 7). At 20 DPI we observe the recruitment of F4/80+ macrophages, Ly6G+ neutrophils, and EPX+ eosinophils to the immature granulomas. By 60 DPI, there is a dramatic expansion of CD68+ F4/80-macrophages, Ly6G+ neutrophils, and accumulation of CD4+ cells at the vasculature. From 90-180 DPI, the immature granulomas condense with decreases in CD68+ F4/80-macrophages, aggregation of Ly6G+ neutrophils in the granuloma core, and increases in CD4+ T cells in the granuloma lymphocyte layer. Mature granulomas with a core of F4/80+ macrophages and Ly6G+ neutrophils with engulfed *C. neoformans* cells surrounded by CD4+ T cells are observed starting at 180 DPI.

**Figure 7:**
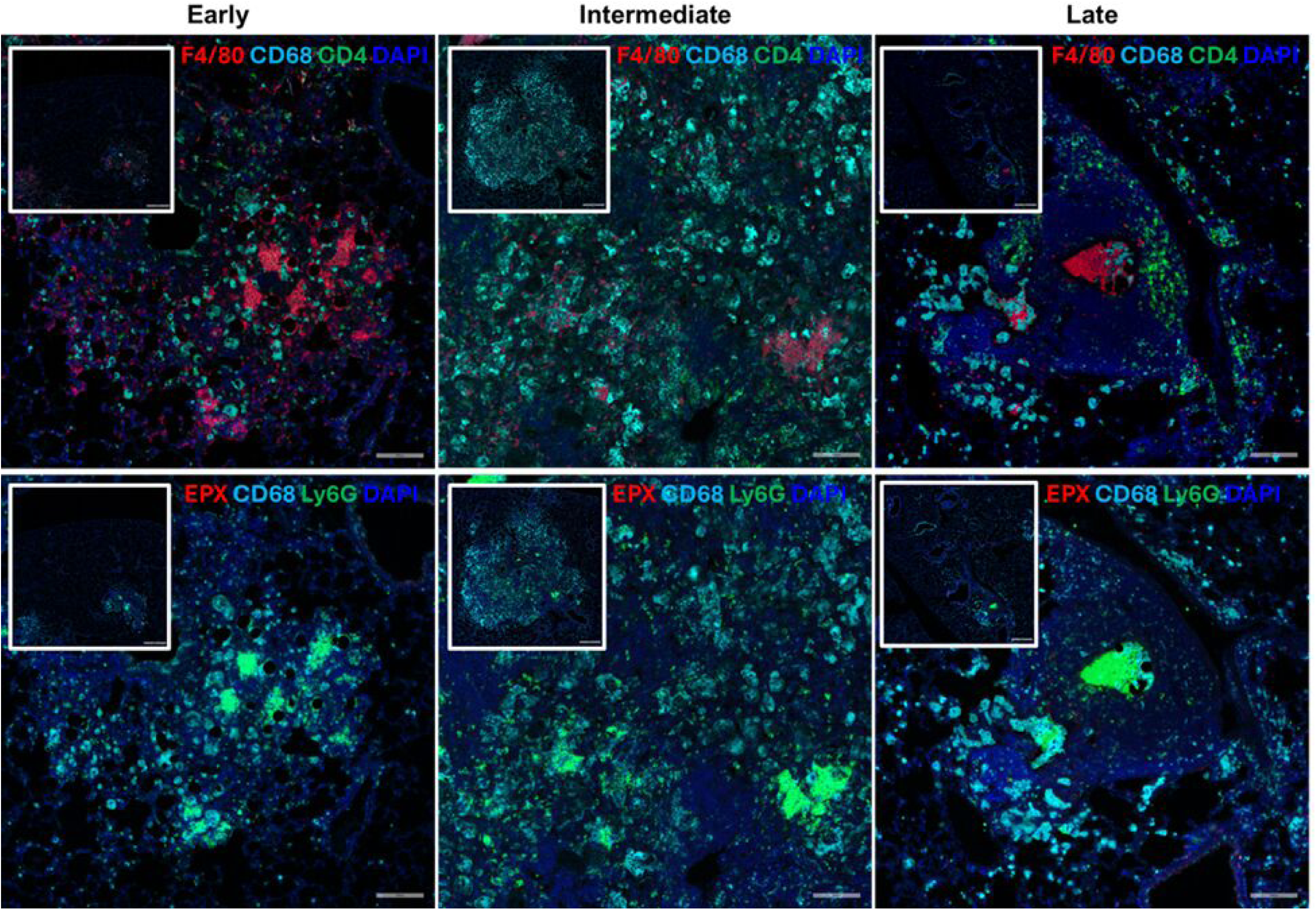
C3HeB/FeJ mice form non-necrotic pulmonary granulomas composed primarily of CD68+ phagocytes, Ly6G+ neutrophils, and CD4+ T cells. Lungs were collected from infected C3HeB/FeJ mice and stained with fluorescent markers targeting F4/80 (red), CD4 (green), and CD68 (cyan) or EPX (red), Ly6G (green), and CD68 (cyan). Representative images at each phase of granuloma formation were taken at 20x magnification while insets were taken at 5x magnification.

## DISCUSSION

Gaining deeper knowledge of immunocompetent host responses against latent *C. neoformans* is crucial for understanding how disseminated cryptococcal disease is prevented in a healthy host. Here, we utilized the C3HeB/FeJ mouse background to model latent Cryptococcus pulmonary infection using the clinical isolate UgCl223 (Fig. 8). We discovered that 1) C3HeB/FeJ mice infected with the latent UgCl223 clinical isolate produce pulmonary granulomas and immune responses distinct from C57Bl/6J and Mycobacterium-infected C3HeB/FeJ mice; 2) that Sp140-deficiency alone is not sufficient to replicate the C3HeB/FeJ cryptococcal response as was previously shown for *Mycobacterium tuberculosis* infections; and 3) latent Cryptococcus infected C3HeB/FeJ mice exhibit early, intermediate and late phases of granuloma formation. These phases of granuloma formation are characterized by different immune responses: a type-2 dominant early response with diffuse granulomatous lesions and *C. neoformans* expansion, a type-1/type-17 dominant intermediate response producing immature granulomas with poor fungal control, and a T cell dominant late response characterized by mature granulomas with effective fungal control. Taken together, these findings paint a picture of the dynamic pulmonary immune response deployed during latent *C. neoformans* infection that can guide future work in understanding the mechanisms involved in preventing disseminated cryptococcal disease.

**Figure 8:**
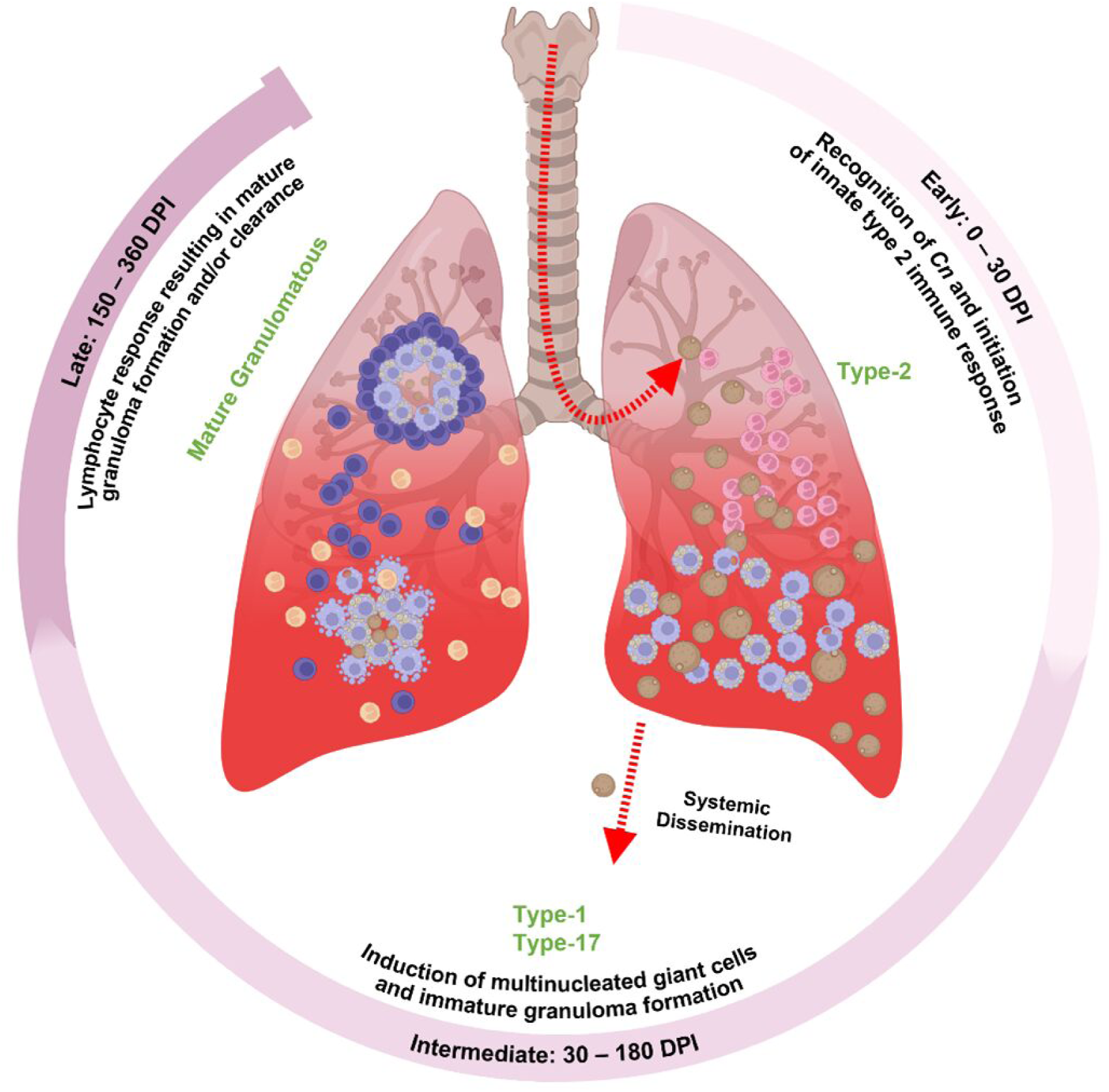
Phases of the pulmonary immune response during latent cryptococcal infection. The histological, pathobiological, and immunological temporal patterns of pulmonary immune responses during latent cryptococcal infection reveal distinct immunological phases. The early phase is characterized by rapidly replicating *C. neoformans* cells, eosinophilia, and IL-4, IL-13, and pro-inflammatory cytokine production. The intermediate phase is characterized by mouse mortality, elevated antigenemia, pro-inflammatory cytokines, and significant increases in interstitial macrophages and CD11b-dendritic cells. The late phase is marked by significant reduction in lung fungal burden and antigenemia, RANTES and IL-10 cytokine abundance, significant increases in both lymphocyte and myeloid cell populations, and mature non-necrotic granuloma formation.

Our studies show that latently infected C3HeB/FeJ mice progress through early, intermediate, and late phases of granuloma formation which are characterized by variation in the immune response (Fig. 8). The early phase is characterized by type-2 responses. Type-2 immune responses in Cryptococcus infection are associated with poor disease outcomes as type-2 cytokine signaling produces immune cells with inferior fungal killing activity ^10,11,20–23^. Indeed, we show here that type-2 responses are ineffectual in preventing cryptococcal expansion as lung colony forming units (CFUs) increased 1000-fold over the first 20 days of the infection. Despite this significant expansion, no mortalities occurred during the early phase of the infection. IL-4, which is highly abundant during the early phase, is important for inducing the formation of MNGCs via the STAT6 signaling mechanism ^24,25^. As MNGCs have historically been shown to be effective at preventing fungal dissemination ^4,7,26^, it is unclear whether the ineffective type-2 immune responses seen during the early phase also initiates a beneficial MNGC response.

The intermediate phase of infection is characterized by a shift to type-1 and type-17 proinflammatory cytokines. Here, we see increased expression of the major type-1 and type-17 cytokines IFNγ and IL-17A, respectively. The protective role of IFNγ in preventing disseminated cryptococcal infection is well established ^20,27,28,21^. In contrast, the role of type-17 immune responses in Cryptococcus infections is unclear, as studies have shown type-17 responses can improve or worsen disease outcomes ^20,29,30^. Enhanced type-17 lymphocyte responses are associated with detrimental neutrophil recruitment ^31^ while knocking out IL-17A signaling leads to increased fungal dissemination ^32^. Therefore, a balanced type-17 immune response is likely important for producing the most benefit during pulmonary cryptococcal infection. Here, we show that the increased expression of IFNγ and IL-17A coincides with the formation of immature granulomas during the intermediate phase of the infection. These granulomas displayed greater organization than the lesions in the early phase but lacked a defined outer lymphocyte layer and were encased in CD68+ phagocytes. Interestingly, despite the increase in protective cytokines, we observed the highest antigenemia burden and mortality during the intermediate phase, suggesting insufficient control of the pulmonary infection (Fig. 8).

The late phase of the infection is characterized by rapid expansion of pulmonary T cell populations, the formation of mature granulomas, and significant elimination of *C. neoformans* cells. T cells, especially CD4+ T cells, are the prevailing immune cell type that confers protection against cryptococcal disease ^5,20,2,22,4^. Our data supports, this as we observe substantial improvements in lung fungal burden, antigenemia, and lung tissue architecture associated with a significant increase in T cell populations in the late phase of granuloma formation. We previously showed that the latent infection model produces a protective type-1 helper T cell (Th1) effector response ^5^. Here, we expanded our analysis of T cell subsets and observed a significant expansion of CD4+ and CD8+ effector, tissue resident memory (Trm), and central memory (Tcm) subsets during the late phase of infection. While significant expansion of effector T cells is seen by 150 DPI, Trm and Tcm populations do not significantly increase until 360 DPI, suggesting the need for additional cellular signaling at these late phases of granuloma formation. In fact, CD11b-DCs, neutrophils, and natural killer cells also significantly increased during the late phase, and we observed mature granulomas with CD4+ T cell rims surrounding neutrophils and phagocytes only at this late stage of the infection. Indeed, it is likely that the late phase experiences T cell-guided myeloid recruitment and modulation that produces protective cellular responses within granulomas. Our findings support that effective control of latent Cryptococcus infection requires T cell expansion but reveal that individual T cell subsets expand differentially rather than as a collective unit, highlighting the need for further investigation into T cell activation and maturation during latent infection.

We found that the cellular immune profiles of the early and intermediate phases of latent Cryptococcus infection are similar to those observed in lethal KN99α infection except in the abundance of CD11b+ dendritic cells (DCs). We observed greater CD11b+ DC recruitment during the early phase of the latent infection than during lethal infection. This is notable as CD11b+ DCs are a heterogenous group of antigen-presenting cells shown to promote type-2 helper T cell (Th2) responses in Cryptococcus infection ^33,23,34^. The binding of *C. neoformans* surface chitin modulates CD11b+ DC activity to promote maladaptive Th2 responses, contributing to KN99α lethality. Type-2 responses dominate the early phase of latent infection, and CD11b+ DCs likely contribute to this response profile. However, our findings reveal that greater CD11b+ DCs abundance does not lead to greater mortality during latent infection. This may point toward a previously unrecognized function of pulmonary CD11b+ DC activity in Cryptococcus infection, such as immunoregulation ^35^, or differences in surface antigen recognition. Overall, we show that lethal and latent Cryptococcal infections share similar initial immune response profiles but differ substantially in their progression toward a beneficial lymphocyte-dominated response.

Our histological analysis reveals that localization of cells during granuloma formation is both temporally and spatially dependent. We determined that the maturation of granulomas in latent Cryptococcus infection consists of 1) early activation of CD68+ MNGCs which phagocytose cryptococcal cells, 2) migration of CD68+ cells to the outer edges of the granuloma while CD4+ T cells enter the core of the immature granuloma, and 3) condensing of the granuloma until CD4+ T cells surround a F4/80+ and Ly6G+ core that also contains the fungal cells. Our timeline shows that CD68+ phagocytes act on the frontline while CD4+ T cells move into areas where fungal cells have already been engaged but not killed. The condensing of the granuloma likely represents the removal of CD68+ phagocytes that successfully cleared fungal cells, while the recruited CD4+ T cells remain surrounding areas of ongoing infection. The lack of CD68+ cells in the core of these mature granulomas suggests that CD68+ phagocytes may be preferred for fungal killing over fungal containment. Our observation of cells singularly positive for either CD68 or F4/80 was unexpected. We report F4/80+ CD68-phagocytes localizing with Ly6G+ neutrophils in granuloma cores during various stages of granulomatous maturation in the early, intermediate, and late phases. In contrast, F4/80-CD68+ phagocytes are primarily observed at the granuloma edge during the intermediate phase and decrease in abundance as granulomas mature and condense. Interestingly, both markers are pan-markers for mouse macrophages ^36^. Previous studies show that macrophage expression of F4/80 is dependent on both the stage of monocyte differentiation and cytokine stimulation. CD68+ monocyte precursors express low levels of F4/80, with increasing F4/80 levels during differentiation ^37^. Likewise, macrophage F4/80 expression decreases in response to IFNγ stimulation during infection ^38^. Here we showed that pulmonary IFNγ expression peaks during the intermediate phase, which coincides with the recruitment of F4/80-CD68+ phagocytes to the granuloma. Therefore, our immunofluorescence studies likely depict either CD68+ monocytes trafficking to granulomas to combat Cryptococcus infection or IFNγ-stimulated tissue-resident macrophages that downregulate F4/80 during the intermediate phase. While F4/80 is expressed primarily on macrophages, it can also be found on other tissue-resident antigen-presenting cells such as dendritic cells (DCs) ^39,40^. Ultimately, our histological analysis reveals that granuloma formation is dependent on the outcomes of CD68+ phagocyte interactions with *C. neoformans* cells, and that differences in CD68 and F4/80 positivity predicts localization within the granuloma at varying phases.

Our findings highlight that the role of Sp140 in the C3HeB/FeJ immune response to latent Cryptococcus infection differs from that seen in Mycobacterium infection. C3HeB/FeJ mice are known to contain a unique allele encoding the super susceptibility to tuberculosis 1 (sst1) locus, a region on chromosome 1 that influences the immune response during Mycobacterium infection ^13,15,41^. The unique sst1S allele in C3HeB/FeJ mice contains deficiencies in immune regulators such as Sp140, a negative IFN1 regulating gene expressed in all immune cells ^16,17,41^. Deficiencies in Sp140 lead to increased expression of IFN1 and other interferon-stimulated genes. Sp140 deficiency replicates the C3HeB/FeJ immune phenotype in C57Bl/6J mice during Mycobacterium infection. We show here that latently infected C3HeB/FeJ mice do not produce IFN1 while lethally infected mice do and that Sp140 deficiency in C57Bl/6J does not significantly impact the immune response to latent Cryptococcus infection. The role of IFN1 in cryptococcal infections is unclear. While IFN1 was found to be present during early infection and induces the release of other beneficial cytokines ^42^, blocking IFN1 signaling was also shown to improve fungal clearance and IFNγ production ^43^. Here, our findings support that IFN1 is unnecessary for effective fungal control and that Sp140 does not play a relevant role in the immune phenotype generated during C3HeB/FeJ latent cryptococcal infection.

Previous studies modeling the C3HeB/FeJ immune response during Mycobacterium infection showed that neutrophil, T cell, and type-1/17 responses are responsible for necrotic granuloma formation and disease outcomes in both *M. avium* and *M. tuberculosis* infections ^44–46^. We show that latent Cryptococcus infection also induces neutrophil and type-1/17 responses during the intermediate phase of granuloma formation, but that the granulomas formed are non-necrotic. However, like in the Mycobacterium models, neutrophil dominance in Cryptococcus infection increased the likelihood of mortality, suggesting that neutrophil presence within granulomas reduces their effectiveness. Additionally, the similarities between elicited type-1/17 cytokine responses in both models may point toward predisposition in C3HeB/FeJ mice for these immune pathways.

Ultimately, the findings of this study provide deeper insight into the previously unknown immunological characteristics of latent pulmonary Cryptococcus infection and granuloma formation. We show that C3HeB/FeJ immune responses to latent Cryptococcus infection are dynamic and transition between three phases of granuloma formation with different genetic influencers compared to Mycobacterium infections. The role these phases play in establishing an effective defense against cryptococcal disease and the influence of the pathogen in modulating granuloma development should be investigated further.

## ACKNOWLEDGMENTS

We thank Yongjuan Xia in the Pathology Services Core (PSC) at the University of North Carolina – Chapel Hill for expert technical assistance with Histopathology and Digital Pathology. The PSC is supported in part by an NCI Center Core Support Grant (P30CA016086). We also thank the University of Minnesota Flow Cytometry Resource (UFCR) for technical support and assistance.

## ETHICS STATEMENT

The animal study was reviewed and approved by the University of Minnesota Institutional Animal Care and Use Committee.

## AUTHOR CONTRIBUTIONS

JJB, MD, and KN conceived and designed the experiments. JJB, MD, JMY, and IM performed the experiments. JJB, MD, IM, and KN analyzed the data. GC, HMA, DM, and KN contributed reagents, materials, and analysis tools. JJB, MD, and KN wrote the paper. All authors contributed to the article and approved the submitted version.

## REFERENCES

1. Ristow, L.C., and Davis, J.M. (2021). The granuloma in cryptococcal disease. PLOS Pathog. 17, e1009342. 10.1371/journal.ppat.1009342.

2. Rajasingham, R., Govender, N.P., Jordan, A., Loyse, A., Shroufi, A., Denning, D.W., Meya, D.B., Chiller, T.M., and Boulware, D.R. (2022). The global burden of HIV-associated cryptococcal infection in adults in 2020: a modelling analysis. Lancet Infect. Dis. 22, 1748–1755. 10.1016/S1473-3099(22)00499-6.

3. McClelland, E.E., Hobbs, L.M., Rivera, J., Casadevall, A., Potts, W.K., Smith, J.M., and Ory, J.J. (2013). The role of host gender in the pathogenesis of *Cryptococcus neoformans* infections. PLoS ONE 8, e63632. 10.1371/journal.pone.0063632.

4. Hill, J.O. (1992). CD4+ T cells cause multinucleated giant cells to form around *Cryptococcus neoformans* and confine the yeast within the primary site of infection in the respiratory tract. J. Exp. Med. 175, 1685–1695. 10.1084/jem.175.6.1685.

5. Ding, M., Smith, K.D., Wiesner, D.L., Nielsen, J.N., Jackson, K.M., and Nielsen, K. (2022). Use of clinical isolates to establish criteria for a mouse model of latent *Cryptococcus neoformans* infection. Front. Cell. Infect. Microbiol. 11.

6. McDonnell, J.M., and Hutchins, G.M. (1985). Pulmonary cryptococcosis. Hum. Pathol. 16, 121–128. 10.1016/S0046-8177(85)80060-5.

7. Shibuya, K., Hirata, A., Omuta, J., Sugamata, M., Katori, S., Saito, N., Murata, N., Morita, A., Takahashi, K., Hasegawa, C., et al. (2005). Granuloma and cryptococcosis. J. Infect. Chemother. 11, 115–122. 10.1007/s10156-005-0387-X.

8. Baker, R.D., and Haugen, R.K. (1955). Tissue changes and tissue diagnosis in cryptococcosis. A study of 26 cases. Am. J. Clin. Pathol. 25.

9. Mitchell, D.H., Sorrell, T.C., Allworth, A.M., Heath, C.H., McGregor, A.R., Papanaoum, K., Richards, M.J., and Gottlieb, T. (1995). Cryptococcal disease of the CNS in immunocompetent hosts: Influence of cryptococcal variety on clinical manifestations and outcome. Clin. Infect. Dis. 20, 611–616. 10.1093/clinids/20.3.611.

10. Coelho, C., Bocca, A.L., and Casadevall, A. (2014). The intracellular life of *Cryptococcus neoformans*. Annu. Rev. Pathol. Mech. Dis. 9, 219–238. 10.1146/annurev-pathol-012513-104653.

11. Mukaremera, L., McDonald, T.R., Nielsen, J.N., Molenaar, C.J., Akampurira, A., Schutz, C., Taseera, K., Muzoora, C., Meintjes, G., Meya, D.B., et al. (2019). The mouse inhalation model of *Cryptococcus neoformans* infection recapitulates strain virulence in humans and shows that closely related strains can possess differential virulence. Infect. Immun. 87, e00046–19. 10.1128/IAI.00046-19.

12. Specht, C.A., Lee, C.K., Huang, H., Tipper, D.J., Shen, Z.T., Lodge, J.K., Leszyk, J., Ostroff, G.R., and Levitz, S.M. (2015). Protection against experimental cryptococcosis following vaccination with glucan particles containing Cryptococcus alkaline extracts. mBio 6. 10.1128/mBio.01905-15.

13. Kramnik, I. (2008). Genetic dissection of host resistance to *Mycobacterium tuberculosis*: The sst1 locus and the Ipr1 gene. In Current Topics in Microbiology and Immunology (Springer Berlin Heidelberg), pp. 123–148. 10.1007/978-3-540-75203-5_6.

14. Lanoix, J.-P., Lenaerts, A.J., and Nuermberger, E.L. (2015). Heterogeneous disease progression and treatment response in a C3HeB/FeJ mouse model of tuberculosis. Dis. Model. Mech. 8, 603–610. 10.1242/dmm.019513.

15. Pichugin, A.V., Yan, B.-S., Sloutsky, A., Kobzik, L., and Kramnik, I. (2009). Dominant role of the sst1 locus in pathogenesis of necrotizing lung granulomas during chronic tuberculosis infection and reactivation in genetically resistant hosts. Am. J. Pathol. 174, 2190–2201. 10.2353/ajpath.2009.081075.

16. Fraschilla, I., Amatullah, H., Rahman, R.-U., and Jeffrey, K.L. (2022). Immune chromatin reader SP140 regulates microbiota and risk for inflammatory bowel disease. Cell Host Microbe 30, 1370–1381.e5. 10.1016/j.chom.2022.08.018.

17. Ji, D.X., Witt, K.C., Kotov, D.I., Margolis, S.R., Louie, A., Chevée, V., Chen, K.J., Gaidt, M., Dhaliwal, H.S., Lee, A.Y., et al. (2021). Role of the transcriptional regulator SP140 in resistance to bacterial infections via repression of type I interferons. eLife 10. 10.7554/elife.67290.

18. Nielsen, K., Cox, G.M., Wang, P., Toffaletti, D.L., Perfect, J.R., and Heitman, J. (2003). Sexual cycle of *Cryptococcus neoformans var. grubii* and virulence of congenic a and α isolates. Infect. Immun. 71, 4831–4841. 10.1128/iai.71.9.4831-4841.2003.

19. Vidal, J.E., and Boulware, D.R. (2015). Lateral flow assay for cryptococcal antigen: An important advance to improve the continuum of HIV care and reduce cryptococcal meningitis-related mortality. Rev. Inst. Med. Trop. São Paulo 57, 38–45. 10.1590/S0036-46652015000700008.

20. Mukaremera, L., and Nielsen, K. (2017). Adaptive immunity to *Cryptococcus neoformans* infections. J. Fungi 3, 64. 10.3390/jof3040064.

21. Hoag, K.A., Lipscomb, M.F., Izzo, A.A., and Street, N.E. (1997). IL-12 and IFN-γ are required for initiating the protective Th1 response to pulmonary cryptococcosis in resistant C.B-17 mice. Am. J. Respir. Cell Mol. Biol. 17, 733–739. 10.1165/ajrcmb.17.6.2879.

22. Lindell, D.M., Ballinger, M.N., Mcdonald, R.A., Toews, G.B., and Huffnagle, G.B. (2006). Diversity of the T-cell response to pulmonary *Cryptococcus neoformans* infection. Infect. Immun. 74, 4538–4548. 10.1128/iai.00080-06.

23. Wiesner, D.L., Specht, C.A., Lee, C.K., Smith, K.D., Mukaremera, L., Lee, S.T., Lee, C.G., Elias, J.A., Nielsen, J.N., Boulware, D.R., et al. (2015). Chitin recognition via chitotriosidase promotes pathologic type-2 helper T cell responses to cryptococcal infection. PLOS Pathog. 11, e1004701. 10.1371/journal.ppat.1004701.

24. Aghbali, A., Rafieyan, S., Mohamed-Khosroshahi, L., Baradaran, B., Shanehbandi, D., and Kouhsoltani, M. (2017). IL-4 induces the formation of multinucleated giant cells and expression of β5 integrin in central giant cell lesion. Med. Oral Patol. Oral Cir. Bucal 22, e1–e6. 10.4317/medoral.20935.

25. Moreno, J.L., Mikhailenko, I., Tondravi, M.M., and Keegan, A.D. (2007). IL-4 promotes the formation of multinucleated giant cells from macrophage precursors by a STAT6-dependent, homotypic mechanism: contribution of E-cadherin. J. Leukoc. Biol. 82, 1542–1553. 10.1189/jlb.0107058.

26. Shibuya, K., Coulson, W.E., Wollman, J.S., Wakayama, M., Ando, T., Oharaseki, T., Takahashi, K., and Naoe, S. (2001). Histopathology of cryptococcosis and other fungal infections in patients with acquired immunodeficiency syndrome. Int. J. Infect. Dis. 5, 78–85. 10.1016/S1201-9712(01)90030-X.

27. Leopold Wager, C.M., Hole, C.R., Wozniak, K.L., and Wormley, F.L. (2016). Cryptococcus and phagocytes: Complex interactions that influence disease outcome. Front. Microbiol. 7.

28. Davis, M.J., Tsang, T.M., Qiu, Y., Dayrit, J.K., Freij, J.B., Huffnagle, G.B., and Olszewski, M.A. (2013). Macrophage M1/M2 polarization dynamically adapts to changes in cytokine microenvironments in *Cryptococcus neoformans* infection. mBio 4, e00264–13. 10.1128/mBio.00264-13.

29. Wormley, F.L., Perfect, J.R., Steele, C., and Cox, G.M. (2007). Protection against cryptococcosis by using a murine gamma interferon-producing *Cryptococcus neoformans* strain. Infect. Immun. 75, 1453–1462. 10.1128/IAI.00274-06.

30. Zelante, T., De Luca, A., D’ Angelo, C., Moretti, S., and Romani, L. (2009). IL-17/Th17 in anti-fungal immunity: What’s new? Eur. J. Immunol. 39, 645–648. 10.1002/eji.200839102.

31. Wiesner, D.L., Smith, K.D., Kashem, S.W., Bohjanen, P.R., and Nielsen, K. (2017). Different lymphocyte populations direct dichotomous eosinophil or neutrophil responses to pulmonary cryptococcus infection. J. Immunol. Baltim. Md 1950 198, 1627–1637. 10.4049/jimmunol.1600821.

32. Trevijano-Contador, N., Roselletti, E., García-Rodas, R., Vecchiarelli, A., and Zaragoza, Ó. (2022). Role of IL-17 in morphogenesis and dissemination of *Cryptococcus neoformans* during murine infection. Microorganisms 10, 373. 10.3390/microorganisms10020373.

33. Mayer, J.U., Demiri, M., Agace, W.W., MacDonald, A.S., Svensson-Frej, M., and Milling, S.W. (2017). Different populations of CD11b+ dendritic cells drive Th2 responses in the small intestine and colon. Nat. Commun. 8, 15820. 10.1038/ncomms15820.

34. Izumi, G., Nakano, H., Nakano, K., Whitehead, G.S., Grimm, S.A., Fessler, M.B., Karmaus, P.W., and Cook, D.N. (2021). CD11b+ lung dendritic cells at different stages of maturation induce Th17 or Th2 differentiation. Nat. Commun. 12, 5029. 10.1038/s41467-021-25307-x.

35. Li, H., Zhang, G.-X., Chen, Y., Xu, H., Fitzgerald, D.C., Zhao, Z., and Rostami, A. (2008). CD11c+ CD11b+ dendritic cells play an important role in intravenous tolerance and the suppression of experimental autoimmune encephalomyelitis. J. Immunol. Baltim. Md 1950 181, 2483–2493.

36. Wei, Q., Deng, Y., Yang, Q., Zhan, A., and Wang, L. (2023). The markers to delineate different phenotypes of macrophages related to metabolic disorders. Front. Immunol. 14, 1084636. 10.3389/fimmu.2023.1084636.

37. Lee, Y.-S., Kim, M.-H., Yi, H.-S., Kim, S.Y., Kim, H.-H., Kim, J.H., Yeon, J.E., Byun, K.S., Byun, J.-S., and Jeong, W.-I. (2018). CX3CR1 differentiates F4/80low monocytes into pro-inflammatory F4/80high macrophages in the liver. Sci. Rep. 8, 15076. 10.1038/s41598-018-33440-9.

38. Ezekowitz, A., and Gordon, S. (1982). Down-regulation of mannosyl receptor-mediated endocytosis and antigen F4/80 in bacillus calmette-guerin-activated mouse macrophages. Role of T lymphocytes and lymphokines. J. Exp. Med. 155, 1623–1637.

39. Sun, X., Jones, H.P., Dobbs, N., Bodhankar, S., and Simecka, J.W. (2013). Dendritic cells are the major antigen presenting cells in inflammatory lesions of murine Mycoplasma respiratory disease. PLoS ONE 8, e55984. 10.1371/journal.pone.0055984.

40. Vermaelen, K., and Pauwels, R. (2004). Accurate and simple discrimination of mouse pulmonary dendritic cell and macrophage populations by flow cytometry: Methodology and new insights. Cytometry A 61A, 170–177. 10.1002/cyto.a.20064.

41. He, X., Berland, R., Mekasha, S., Christensen, T.G., Alroy, J., Kramnik, I., and Ingalls, R.R. (2013). The sst1 resistance locus regulates evasion of type I interferon signaling by *Chlamydia pneumoniae* as a disease tolerance mechanism. PLOS Pathog. 9, e1003569. 10.1371/journal.ppat.1003569.

42. Qin, H.-J., Feng, Q.-M., Fang, Y., and Shen, L. (2014). Type-I interferon secretion in the acute phase promotes *Cryptococcus neoformans* infection-induced Th17 cell polarization in vitro. Exp. Ther. Med. 7, 869–872. 10.3892/etm.2014.1517.

43. Sato, K., Yamamoto, H., Nomura, T., Matsumoto, I., Miyasaka, T., Zong, T., Kanno, E., Uno, K., Ishii, K., and Kawakami, K. (2015). *Cryptococcus neoformans* infection in mice lacking type I interferon signaling leads to increased fungal clearance and IL-4-dependent mucin production in the lungs. PLOS ONE 10, e0138291. 10.1371/journal.pone.0138291.

44. Gern, B.H., Klas, J.M., Foster, K.A., Cohen, S.B., Plumlee, C.R., Duffy, F.J., Neal, M.L., Halima, M., Gustin, A.T., Diercks, A.H., et al. (2024). CD4-mediated immunity shapes neutrophil-driven tuberculous pathology. bioRxiv, 2024.04.12.589315. 10.1101/2024.04.12.589315.

45. Marzo, E., Vilaplana, C., Tapia, G., Diaz, J., Garcia, V., and Cardona, P.-J. (2014). Damaging role of neutrophilic infiltration in a mouse model of progressive tuberculosis. Tuberculosis 94, 55–64. 10.1016/j.tube.2013.09.004.

46. Verma, D., Stapleton, M., Gadwa, J., Vongtongsalee, K., Schenkel, A.R., Chan, E.D., and Ordway, D. (2019). *Mycobacterium avium* infection in a C3HeB/FeJ mouse model. Front. Microbiol. 10, 693. 10.3389/fmicb.2019.00693.

